## Supplementary materials for "Haplotype-aware modeling of *cis*-regulatory effects highlights the gaps remaining in eQTL data"

**Supplementary Figures**

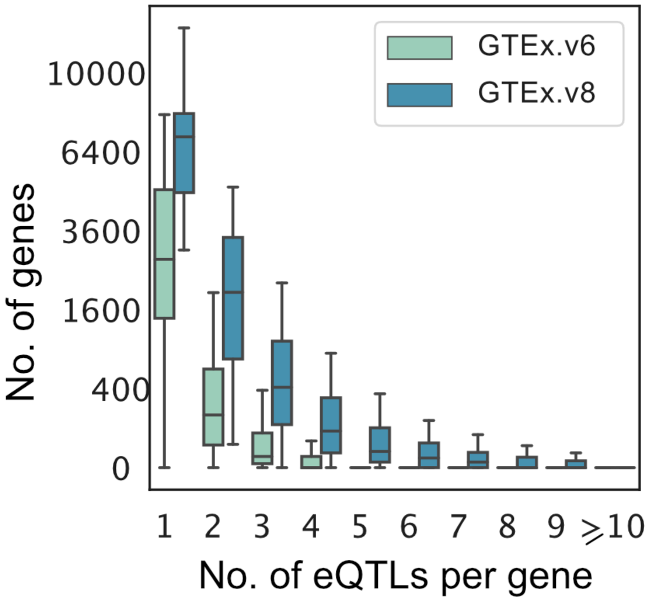


**Supplementary Fig. 1: Plenty of genes are associated with multiple independent regulatory variants**. We compared the quantity of genes as a function of the number of associated eQTLs per gene, across tissues for GTEx v6 and GTEx v8 data. About 70 percent of genes have more than one eQTL in at least one tissue in GTEx v8 data. About 9.4 percent of genes in each tissue have multiple eQTLs in GTEx v6 and this increased to 25.9 percent in GTEx v8 data. There are up to 7 and 16 independent eQTLs per gene for GTEx v6 and v8, respectively.

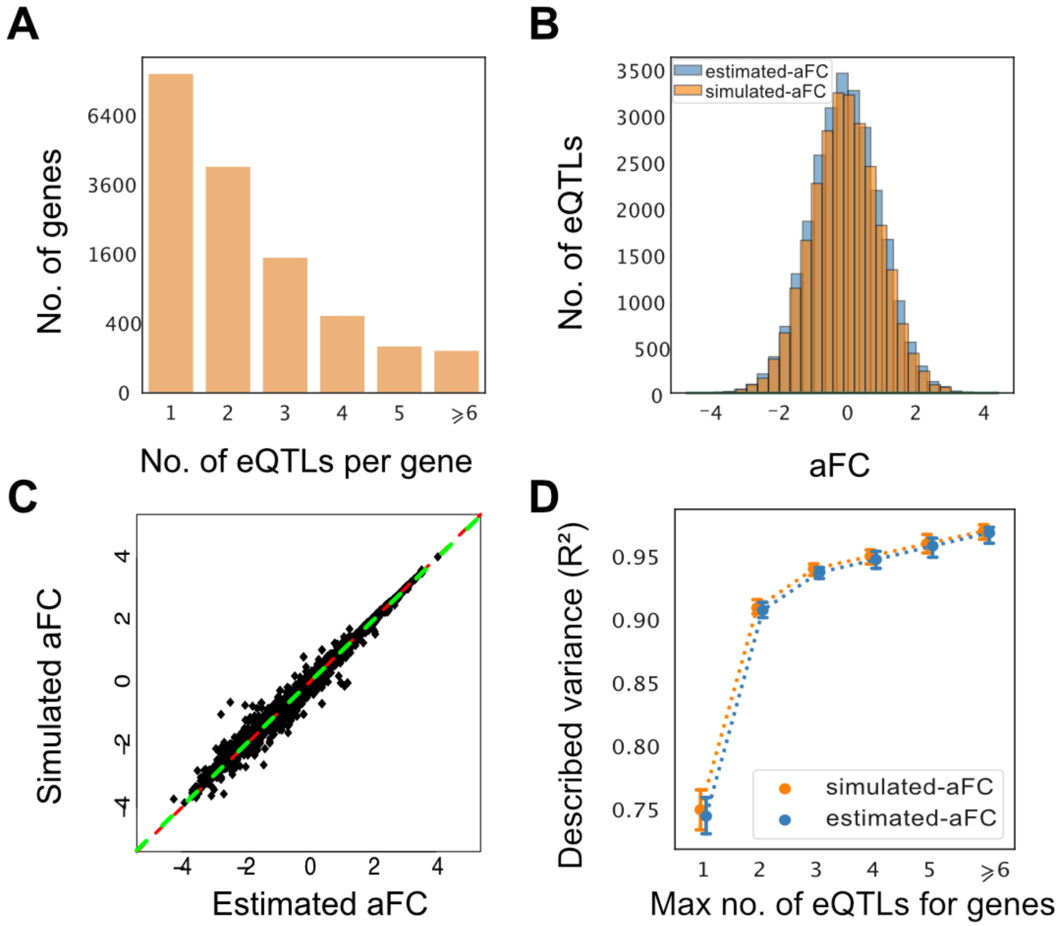


**Supplementary Fig. 2: Estimated effect sizes using the simulated expression data.** A) Distribution of the number of associated eQTLs per gene for 15,167 genes. B-C) Our method provides highly accurate and similar estimates to simulated aFCs. The distribution of estimated aFCs (*norm*[-0.01,$\sigma$=1.01]) is similar to the distribution of simulated aFCs (*norm*[-0.01,$\sigma$=1]) (B). Pearson and Spearman correlation coefficients are both 0.99, and the Deming regression line shown in green is (y= 0.99x+0.00015) and the red line is (y=x) (C). D) Described variance of predicting simulated allelic imbalance with the estimated and simulated effect sizes.

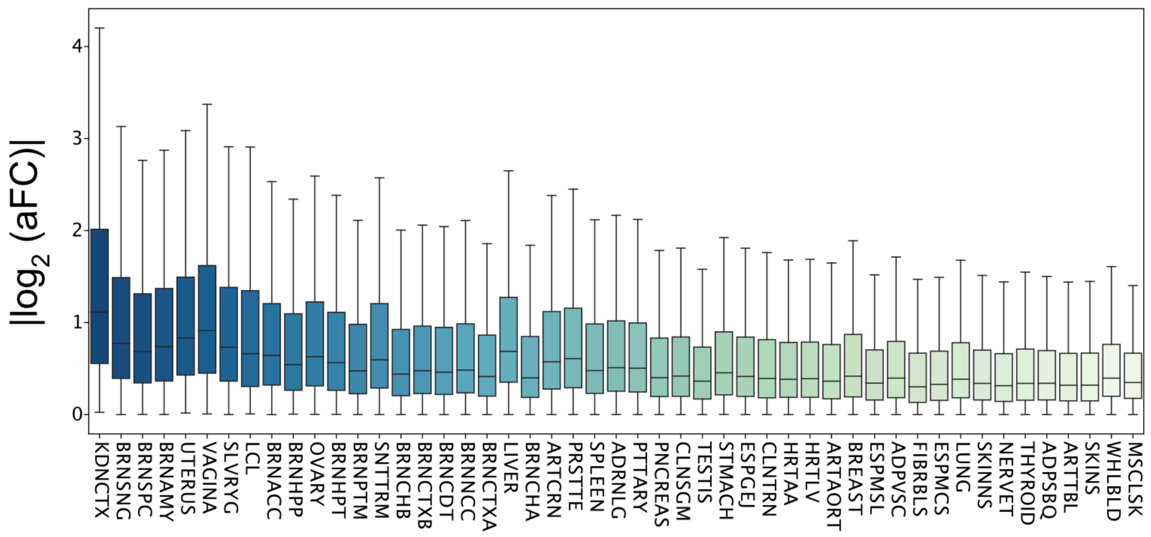


**Supplementary Fig. 3: Distribution of absolute aFC across tissues, sorted from smallest sample size (kidney-cortex, n = 73) to the largest (muscle-skeletal, n = 706).** The distribution of aFCs for *cis*-eQTLs detected in GTEx tissues are dependent on the sample size. There is not sufficient power to detect weak eQTLs for tissues with lower sample sizes.

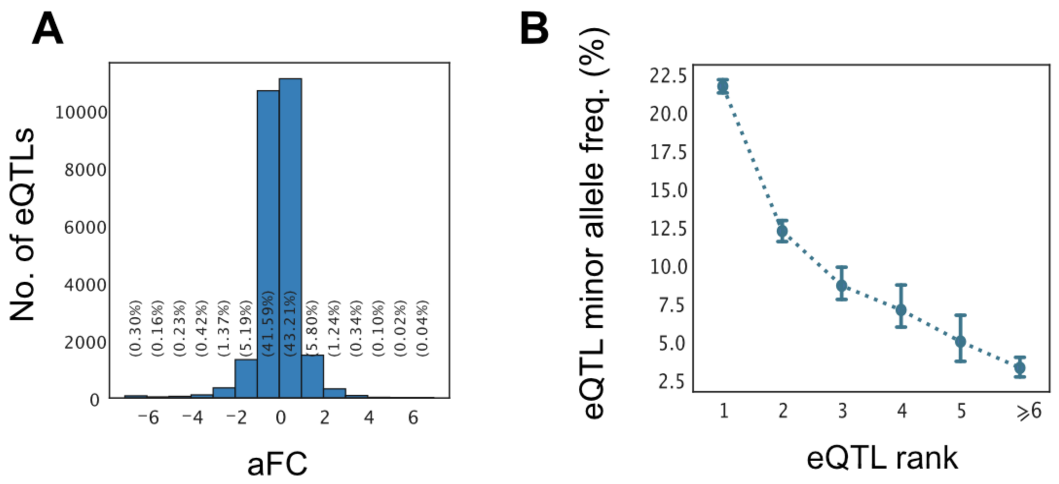


**Supplementary Fig. 4: Empirical properties of estimated aFCs in GTEx v8 adipose subcutaneous tissue.** A) Distribution of estimated aFCs for 25,682 identified eQTLs. B) Minor allele frequency has a decreasing pattern for secondary eQTLs (41.3% of eQTLs are secondary eQTLs). The effect size distribution is affected by the power of eQTL mapping (Fig.2A-B). Error bars represent 95% bootstrap confidence intervals of the median.

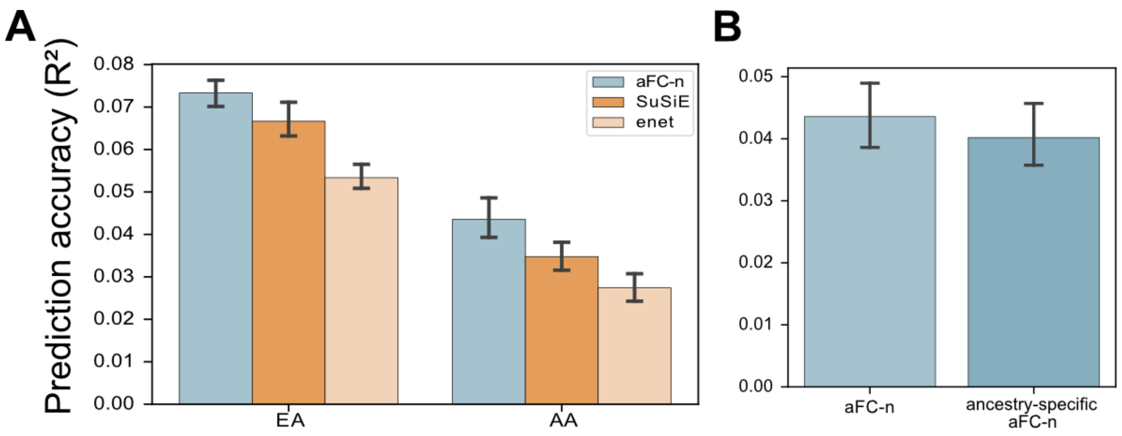


**Supplementary Fig. 5: The expression prediction accuracy is lower among African American (AA) compared to European American (EA) individuals.** The aFCs for independent eQTLs in GTEx v6p data were derived using normalized expressions of adipose subcutaneous samples and tested on unseen GTEx v8 samples. A) Comparison of predicted gene expression with aFC-n, elastic net (enet) and SuSiE for 4,719 genes in common among all models, stratified by self-reported ancestry for the GTEx donors. B) The ancestry-specific aFC-n estimation did not improve the accuracy of the standard aFC-n model on AA expression prediction accuracy. This could be explained as the result of overfitting to the limited training data for the AA population (n=44) and also failing to reflect admixed haplotype structure in African American donors.

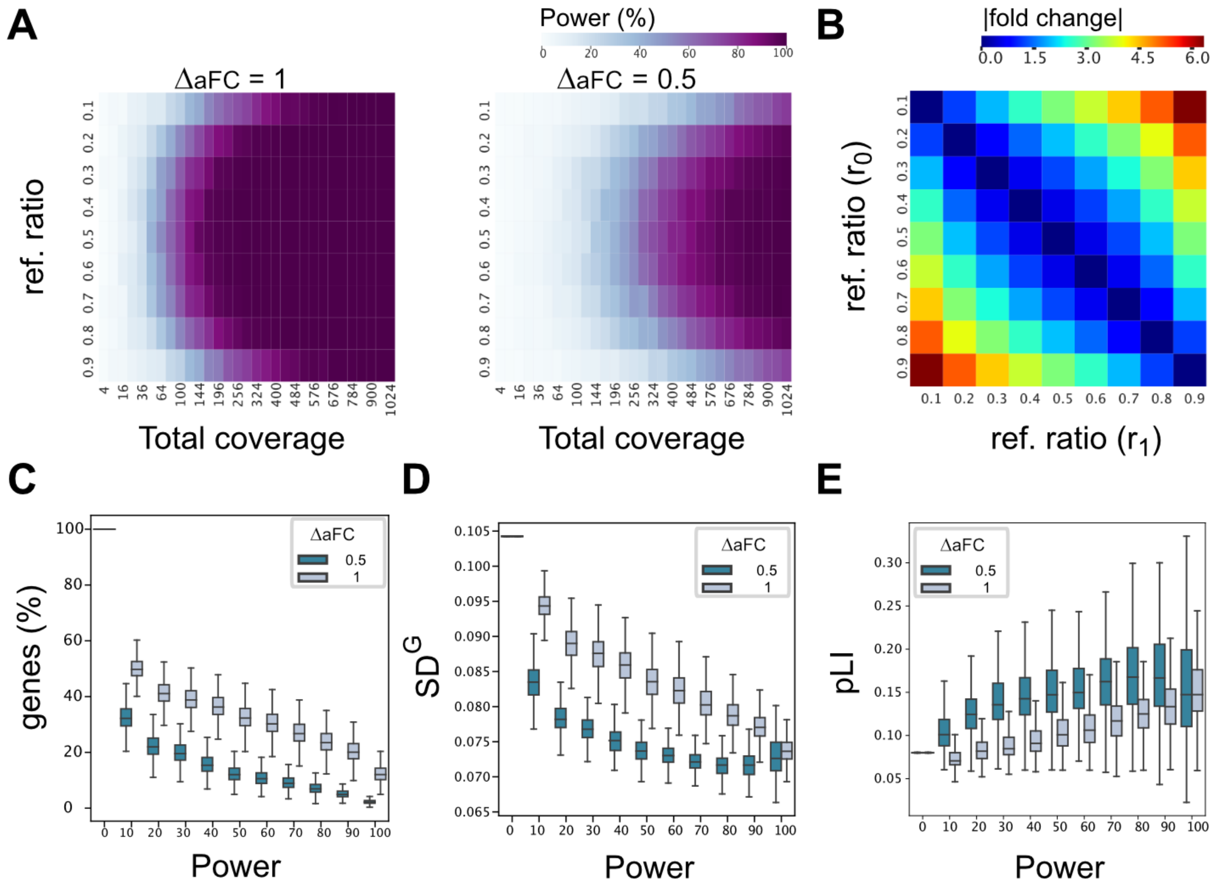


**Supplementary Fig. 6: Power analysis to estimate the fraction of the cases that the current eQTL data does not fully describe ASE signal**. A) Power estimation based on simulation for a set of read counts and reference ratios for the specific fold changes ($\Delta aFC$) 0.5 and 1. In low count cases there is not enough power to confirm the difference between the observed and predicted values, and improvement in power is observed by increase in the expression read counts. B) The absolute fold change between the $\log_2$(aFC)s is illustrated as a difference between the reference ratios. The reference ratio is defined as the logistic function of the log(aFC) using Eq.5. C) The total number of genes at different levels of power for adipose subcutaneous tissue. This indicates that for 6.9 and 23.5 percent (median) of genes in an individual, there is 80 percent power to detect 0.5 and 1 fold changes, respectively. The percentage of genes with an excess allelic imbalance is presented in Fig.4A. D) Median of $SD^G$ estimates across samples as a function of power. Highly expressed genes with high statistical power tend to be also less tolerant to variation. E) Median of pLI (Probability of loss of function intolerance) across samples as a function of power. Highly expressed genes with high statistical power are less tolerant to protein truncating variation.

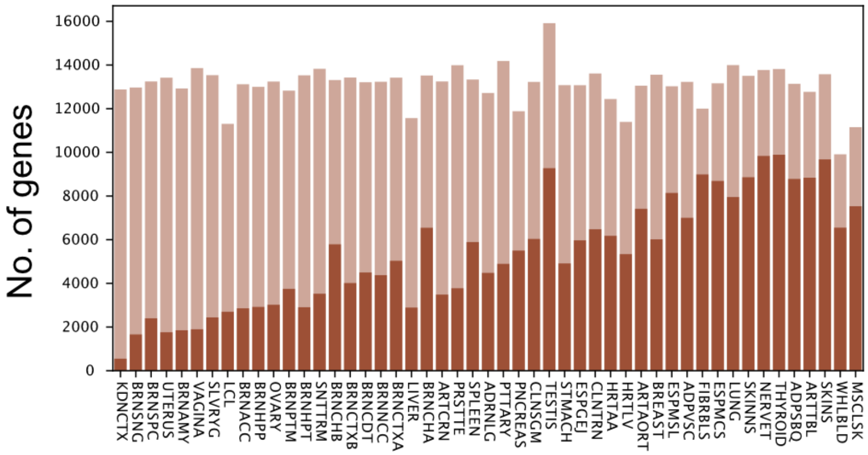


**Supplementary Fig. 7: The number of eGenes grows with the sample size.** Amount of all protein-coding genes with median TPM >1 (shown with light bar) and the number of autosomal eGenes (shown with dark bar) per tissue (Spearman corr. = 0.92).

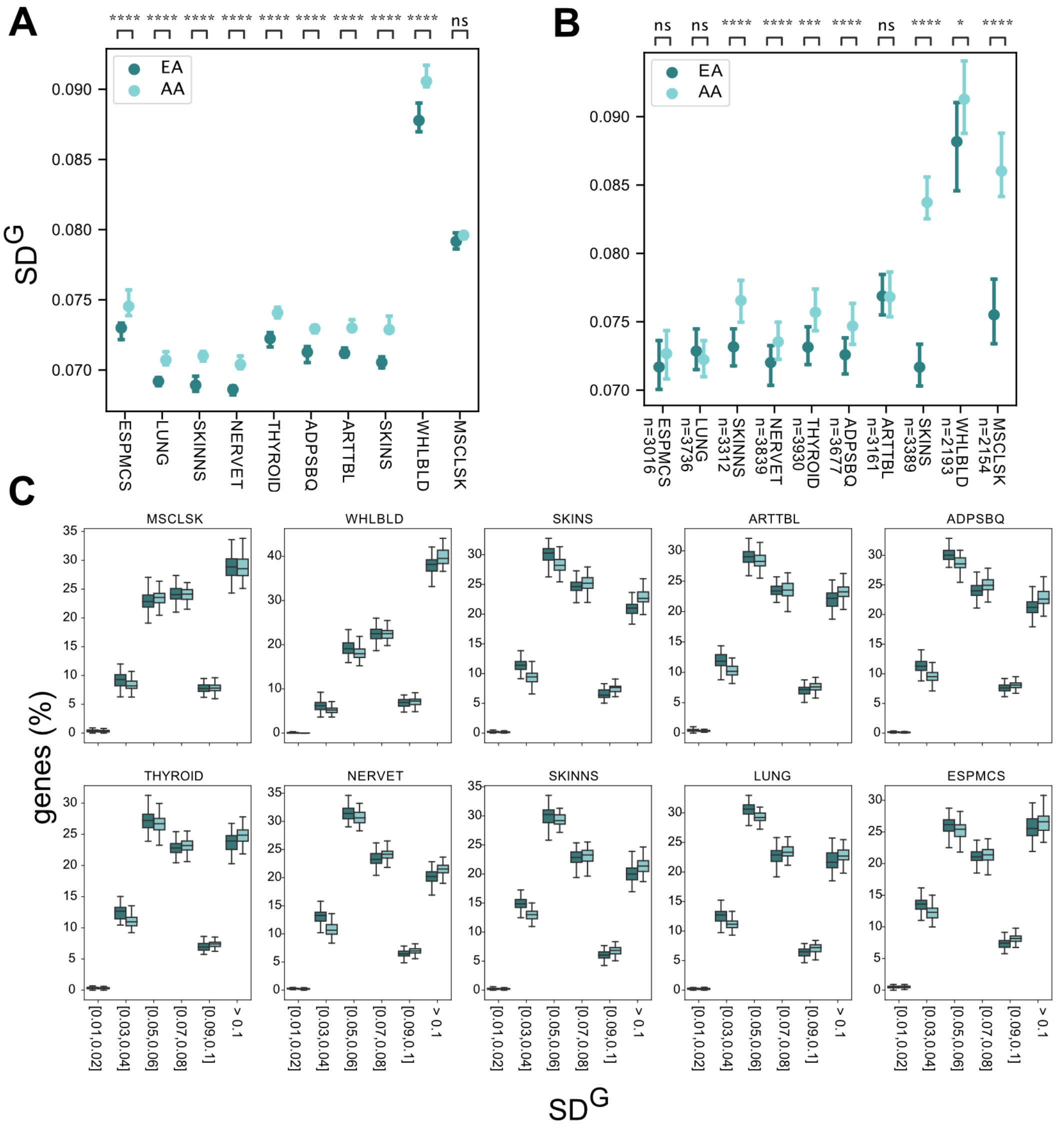


**Supplementary Fig. 8: $SD^G$ estimates across African American and European American populations in top 10 sampled tissues**. A-B) The genetic dosage variation is more variable in the AA population. The median of $SD^G$s across samples (A). The expression variations of the common genes in AA and EA populations present in at least one sample analyzed in (Fig.5C). The number of genes for each tissue is specified in the x-axis. The $SD^G$s are calculated over 60 random samples of each population (B). Two-sided Wilcoxon signed rank test with Bonferroni correction p-value annotation: ns: 5.00E-02 < p <= 1.00; *: 1.00E-02 < p <= 5.00E-02; **: 1.00E-03 < p <= 1.00E-02; ***: 1.00E-04 < p <= 1.00E-03; ****: p <= 1.00E-04. C) Proportion of tested genes as a function of $SD^G$ bins. The x-axis is the ancestry agnostic $SD^G$ capped at 0.1 for ease of viewing. African Americans have a larger proportion of genes with higher variation.

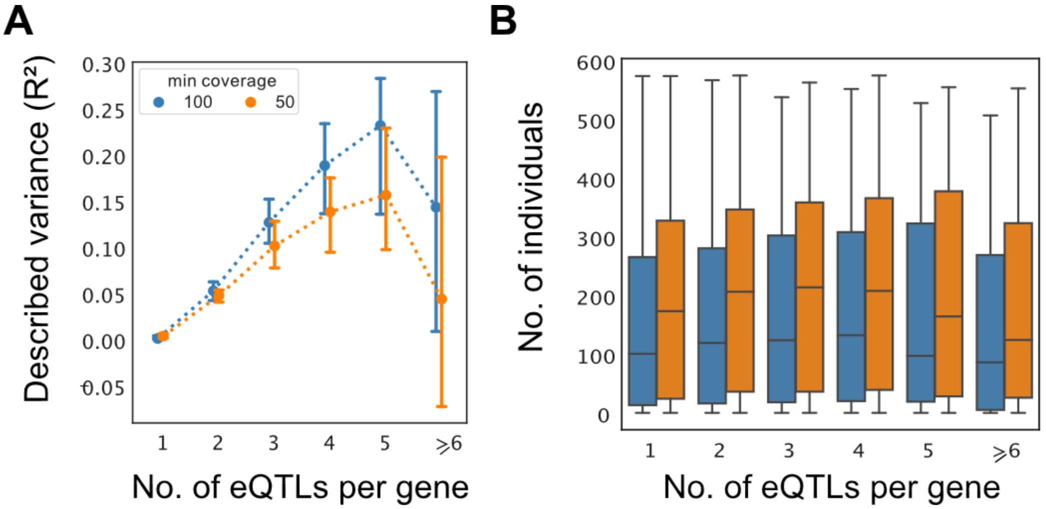


**Supplementary Fig. 9: Described variance and amount of ASE data available at the minimum read coverages of 100 and 50**. A-B) Described variance of predicting allelic imbalance (A), and the number of individuals with ASE data (B) as a function of eQTL counts per gene subject to minimum read coverage of 100 and 50 from adipose subcutaneous tissue. Increasing minimum read coverage from 50 to 100, increased the $R^2$ while missing 1,435 (12.6%) genes. Error bars represent 95% bootstrap confidence intervals of the median.

### Supplementary Tables

**Supplementary Table 1: Tissue abbreviations**

**Supplementary Table 2: Sample size and median number of genes among samples analyzed in Fig.4B.**

**Supplementary Table 3: Sample size and median number of genes among samples analyzed in Fig.5B.**
